# Soil bacterial populations are shaped by recombination and gene-specific selection across a meadow

**DOI:** 10.1101/695478

**Authors:** Alexander Crits-Christoph, Matthew Olm, Spencer Diamond, Keith Bouma-Gregson, Jillian Banfield

## Abstract

Soil microbial diversity is often studied from the perspective of community composition, but less is known about genetic heterogeneity within species and how population structures are affected by dispersal, recombination, and selection. Genomic inferences about population structure can be made using the millions of sequencing reads that are assembled de novo into consensus genomes from metagenomes, as each read pair describes a short genomic sequence from a cell in the population. Here we track genome-wide population genetic variation for 19 highly abundant bacterial species sampled from across a grassland meadow. Genomic nucleotide identity of assembled genomes was significantly associated with local geography for half of the populations studied, and for a majority of populations within-sample nucleotide diversity could often be as high as meadow-wide nucleotide diversity. Genes involved in specialized metabolite biosynthesis and extracellular transport were characterized by elevated genetic diversity in multiple species. Microbial populations displayed varying degrees of homologous recombination and recombinant variants were often detected at 7-36% of loci genome-wide. Within multiple populations we identified genes with unusually high site-specific differentiation of alleles, fewer recombinant events, and lower nucleotide diversity, suggesting recent selective sweeps for gene variants. Taken together, these results indicate that recombination and gene-specific selection commonly shape local soil bacterial genetic variation.

## Introduction

Soil microbial communities play key biogeochemical roles in terrestrial ecosystems (Fierer 2017). Within a single hectare of temperate grassland soil, there can be over 1,000 kg of microbial biomass and corresponding large microbial population sizes (Fierer et al. 2009). However, many of the most abundant soil microorganisms are underrepresented or absent from culture collections and genomic databases, even at the level of class or phylum (Lloyd et al. 2018). Progress has been made in cataloging the diversity of 16S rRNA genes in soils (Thompson et al. 2017), which is useful for understanding microbial community composition, but this technique is incapable of discerning genetic variation within populations. On the other hand, genome-resolved metagenomics studies, in which shotgun sequencing of metagenomic DNA is assembled and binned into draft genomes, have resulted in whole genome characterization of several highly abundant soil bacteria (Hultman et al. 2015; Butterfield et al. 2016; White et al. 2016; Ji et al. 2017; Woodcroft et al. 2018; Diamond et al. 2019).

Sequence variation within genome-resolved metagenomic datasets can also be used to track changes in allele frequencies, and to infer the operation of evolutionary forces of genetic drift, natural selection, and homologous recombination (Whitaker and Banfield 2006; Garud et al. 2019). Homologous recombination, in particular, can vary dramatically in its importance relative to other processes for different microbial species. Analyses of reference genomes have shown that homologous recombination frequently occurs in bacteria populations, both globally and locally (González-Torres et al. 2019; Lin and Kussell 2019; Sakoparnig et al. 2019). For example, certain populations of hotspring Cyanobacteria approach panmixia, where recombination is so frequent that individual cells are unlinked random mixtures of alleles (Rosen et al. 2015). In other species like oceanic Vibrio, recombination is high but large blocks of alleles important for ecological niche differentiation are co-inherited and may remain linked due to selection (Cui et al. 2015) (Shapiro et al. 2012). In soils, Streptomyces flavogriseus isolates were also found to approach a freely recombining panmixia (Doroghazi and Buckley 2010). In contrast, Myxoccocus xanthus isolates recovered from a series of soil samples were distinct but highly clonal, implying recombination between strains was low (Wielgoss et al. 2016). The degree of recombination in most abundant soil bacterial lineages has not been investigated, but it may be widespread, as high cell densities could promote the sharing of genetic material via transformation (Thomas and Nielsen 2005), conjugation (Rocha et al. 2005), or the uptake of extracellular vesicles (Tran and Boedicker 2019).

When recombination rates are low or selection is extremely strong, several clonal strains can compete until one or more beneficial alleles is highly selected for, resulting in a single clonal genotype sweeping to fixation. However, when recombination unlinks gene variants within a population, beneficial alleles can sweep through a population independent of genomic context in a selective sweep (Smith and Haigh 1974) with gene/locus-specific effects. One genome-resolved metagenomic study observed a single clonal sweep over a nine year period for one Chlorobium population in a freshwater lake (Bendall et al. 2016), while most of the other bacterial populations studied possessed genomic loci with unusually few SNPs (single nucleotide polymorphisms), an observation interpreted as evidence for gene-specific selective sweeps. However, there are additional signals within genomic data that could further constrain evolutionary history, as positive selection acting on a genomic locus in a recombining population can leave locus-specific signals relative to the genomic average. These include higher linkage disequilibrium (a strong association between alleles) and different allele frequencies between populations (Krause and Whitaker 2015) (Shapiro et al. 2009). In soil microbial populations, comparative frequencies of gene-specific sweeps versus genome-wide clonal expansions are largely unknown, and it is generally uncertain whether selection or dispersal constraints play a larger role in shaping allele frequencies.

Previously, we conducted a large scale genome-resolved metagenomics study of soils from a grassland meadow in the Angelo Coast Range Reserve in northern California that established a dataset of 896 dereplicated phylogenetically diverse microbial genomes (Diamond et al. 2019). The soil at the site is a sandy loam with pH values in the range of 4.6 to 4.9, and the grassland is dominated by annual Mediterranean grasses and forbs (Suttle et al. 2007; Butterfield et al. 2016). The meadow has been part of a rainfall amendment climate change study ongoing for over 17 years and has also been studied in the context of plant diversity and productivity (Sullivan et al. 2016), invertebrate herbivores and predators (Suttle et al. 2007), fungal communities (Hawkes et al. 2011), soil organic matter (Berhe et al. 2012), metabolomics, and metaproteomics (Butterfield et al. 2016). By analyzing the population genomics of 19 highly abundant bacterial species across this meadow, we found high nucleotide diversity within samples, intrapopulation genetic structure often explained by local spatial scales, varying degrees of homologous recombination for different species, and gene-specific population differentiation partially driven by selection.

## Results and Discussion

### Bacterial allele frequencies within populations are spatially organized across a grassland meadow

Previously, 60 soil samples were collected at depths of 10-20, 20-30, and 30-40 cm from within six 10 m diameter plots (see *Methods*). Samples were collected over a period of three months (**Fig 1a**). Sampling plots were arranged in blocks of two plots, and one of the two plots in each block received extended spring rainfall ((Suttle et al. 2007).

**Figure 1.**
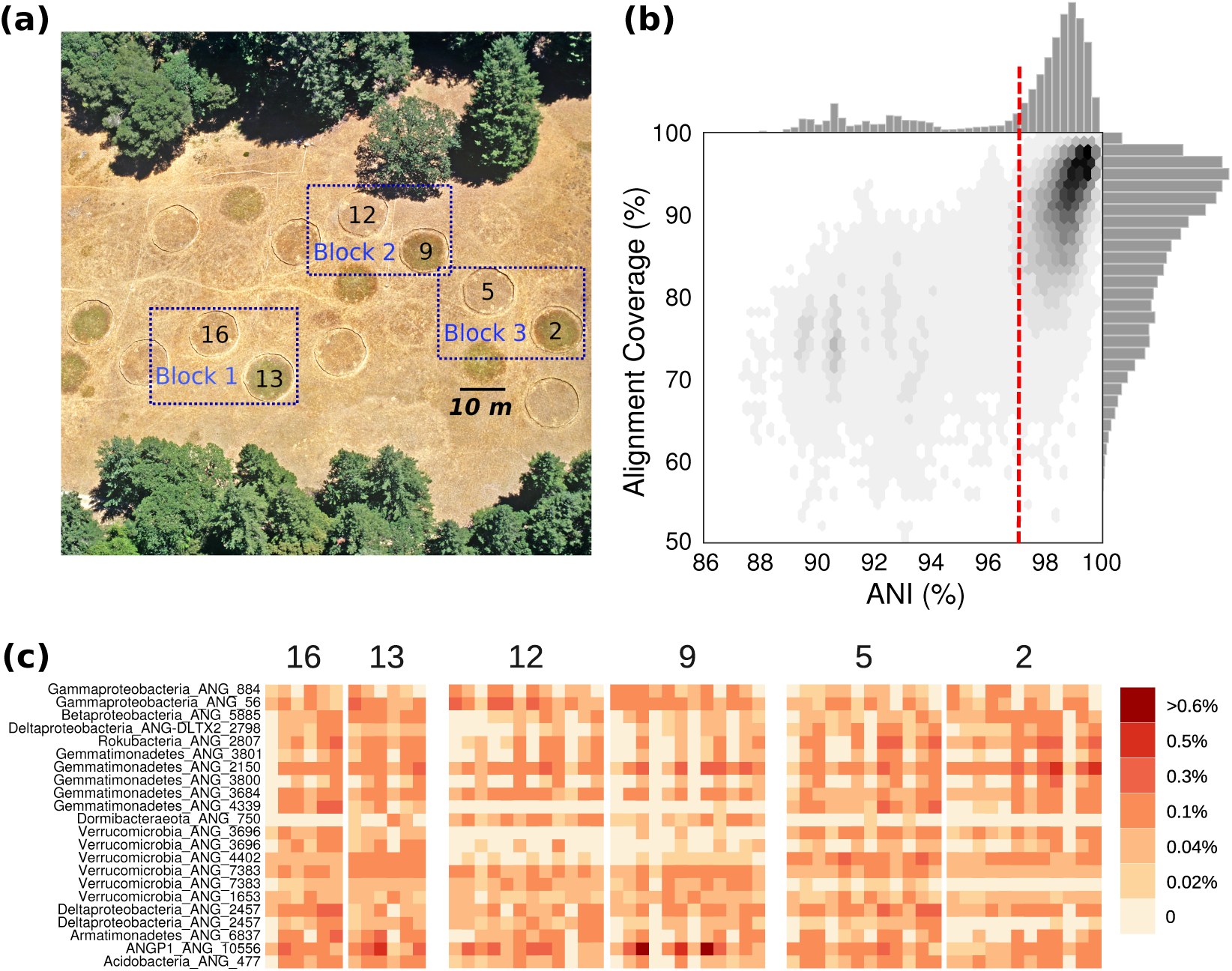
Meadow overview and population relative abundances. **(a)** A bird’s eye view of the grassland meadow located at the Angelo Coast Range Reserve in Mendocino County, California (39° 44’ 17.7” N, 123° 37’ 48.4” W). **(b)** Histograms of average nucleotide identity (ANI) and alignment coverage (an approximation of % shared gene content) values for all-vs-all comparisons between genomes assembled from the meadow soils. **(c)** Relative abundances of the 19 bacterial populations analyzed in this study in each sample, organized by experimental plot where sample was collected.

Starting with the dataset of 3,215 draft quality genomes assembled from samples collected from across the meadow (Diamond et al. 2019), we calculated all pairwise genome-wide average nucleotide identities (ANI) and alignment coverages (roughly analogous to shared gene content; **Fig 1b**). We observed a sharp decrease in pairwise ANI for all of the genomes from the meadow around 96.5-97%, similar to the threshold for bacterial species delineation reported recently (Jain et al. 2018). We observe a more gradual decline in shared gene content, indicating that within species, even within a local environment, shared gene content can often be more variable than average nucleotide identity of conserved regions. We used the 97% ANI cutoff to cluster genomes into groups of species-like populations. Some species-like groups contained dozens of near-complete draft genomes, each assembled from a different sample independently, and we selected species clusters with at least 12 genomes estimated to be >80% complete with <10% contamination for population genetics analysis. The final set comprised 467 genomes from 19 widespread populations (312 genomes are estimated to be >90% complete; **Supplementary Table S1**).

The bacterial species in this set included many commonly reported highly abundant soil bacteria from phyla including Chloroflexi, Acidobacteria, Verrucomicrobia, and Rokubacteria, which are known to be abundant globally in soils (Fierer 2017; Bergmann 2011), but remain understudied. Most of the species in this set were likely novel at the taxonomic rank of class, and one likely represents a novel candidate phylum tentatively designated ANGP1 (Diamond et al. 2019). Based on measurement of the relative abundance of each population across the entire meadow, these bacteria are some of the most abundant species in the soil, although no individual species contributed >1% of the DNA in a sample (**Figure 1c**).

For each of the 19 meadow-wide populations, we tested to see if the proximity of sampling sites within meadow plots predicted genetic similarity of the assembled genomes (PERMANOVA; FDR<=5%; adjusted p<=0.05). Further, we tested if samples collected from the same soil depth were more similar than those collected from different depths. We found that the genetic variation of genomes from 12 of the 19 populations were significantly associated with sampling plot, and that genetic variation within 5 of the 19 populations were significantly associated with sampling depth (**Fig 2a**). Non-metric multidimensional scalings of the nucleotide identity matrices of genomes from each population showed clear associations with both plot of origin and depth (**Fig 2b**). Because the genome assembly from each sample reflects the most abundant sequence variant in each population, this implies that major allele frequencies varied across the meadow. While local spatial heterogeneity has been shown to highly explain microbial community composition in soils (O’Brien et al. 2016), here we demonstrate that there are also spatial patterns within the genetic variation of individual species.

**Figure 2.**
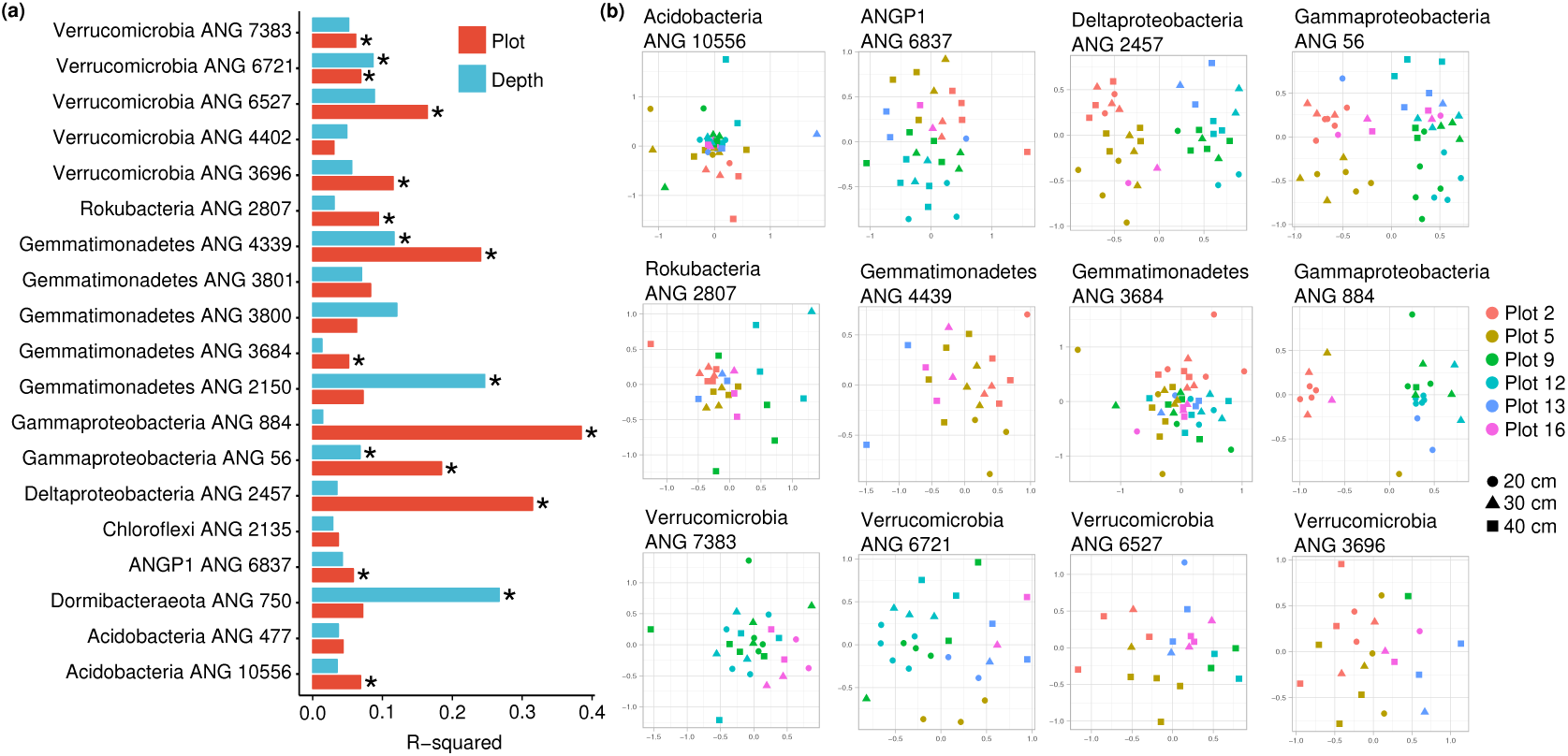
Spatial variation in genetic differences within species. **(a)** The percentage of variation in genetic similarity (ANI) of consensus genomes explained by plot of origin (red) and sampling depth (blue). **(b)** NMDS of genetic dissimilarities between genomes within each species, for all 12 populations for which sampling plot explained a significant fraction of the variation in genetic dissimilarity (PERMANOVA; FDR=5%; p<0.05). There is a single point for each genome independently assembled for a population, and genomes are colored by sample plot of origin and the point shape indicates the sample depth.

### High population nucleotide diversity is observed meadow-wide and within soil samples

Each genome was assembled from >100,000 short reads, and millions of reads could be assigned to each population from all 60 samples. Because microbial populations within the 10 g soil samples used for DNA extraction are comprised of orders of magnitude more cells than were sequenced, most read pairs are likely from unique cells (or DNA molecules) in the population. Each genome assembled from each sample was sequenced at around 10× coverage, but meadow-wide coverages for each population ranged from 224× to 908× (**Supplementary Table S2**).

We implemented stringent requirements for read mapping of read pairs that were at least 96% combined sequence identity to a representative reference for each genome (see Methods). Replicate genomes from each species were used to eliminate possible contaminating contigs within each metagenome-assembled genome assembly, and SNPs were then called at frequencies >5% in each population using reads from all samples, using a simple null model that assumes a false discovery rate <∼10^-6^. Compared to a previously published study on freshwater lake microbial populations (Bendall et al. 2016), there were far more polymorphisms per population (4,721 to 43,225 SNPs / Mbp, **Supplementary Table S2**), despite having a minimum allele frequency cutoff of 5%. Most populations also had higher rates of SNPs / Mbp than a similar analysis of metagenome assembled genomes found for microbial populations in deep-sea hydrothermal vents (Anderson et al. 2017).

To assess the genetic variability within each population across the meadow beyond the SNPs / Mbp metric, we calculated the per-site nucleotide diversity of the sequencing reads (probability that any two sampled reads share the same base at a site) at each locus for that population. Previously, metagenomic studies have used either the average similarity of reads to a reference or the total number of SNPs / Mbp as metrics of genetic diversity (Bendall et al. 2016) (Anderson et al. 2017). We chose to measure nucleotide diversity because (1) it is less sensitive to large changes in coverage (**Fig S1**), (2) it can be calculated both for a single site and averaged over genes or windows, and (3) it considers not only the number of SNPs but also their frequencies in the population. We found a wide range of per-gene nucleotide diversity values for the 19 different populations (**Fig 3a**). Because nucleotide diversity is less sensitive to changes in coverage, we could use it to track how genetic diversity changes with distance across the meadow. We calculated nucleotide diversity in pooled mapped reads for each population sampled from the same location and soil depth, within plot, within block (pairs of plots), and across the entire meadow for each species (**Fig 3a**). Although nucleotide diversity tends to be higher at the meadow scale compared to the sample scale, the genetic diversity within some samples was comparable to that across the entire meadow for many populations, indicating that in some cases high genetic diversity persists within soils even at the centimeter scale. This implies that although there are shifts in allele frequencies across the plots within the meadow, within each 10 g sample of soil many alternative alleles are frequently encountered, and extensive gene variant dispersal occurs across the meadow.

**Figure 3.**
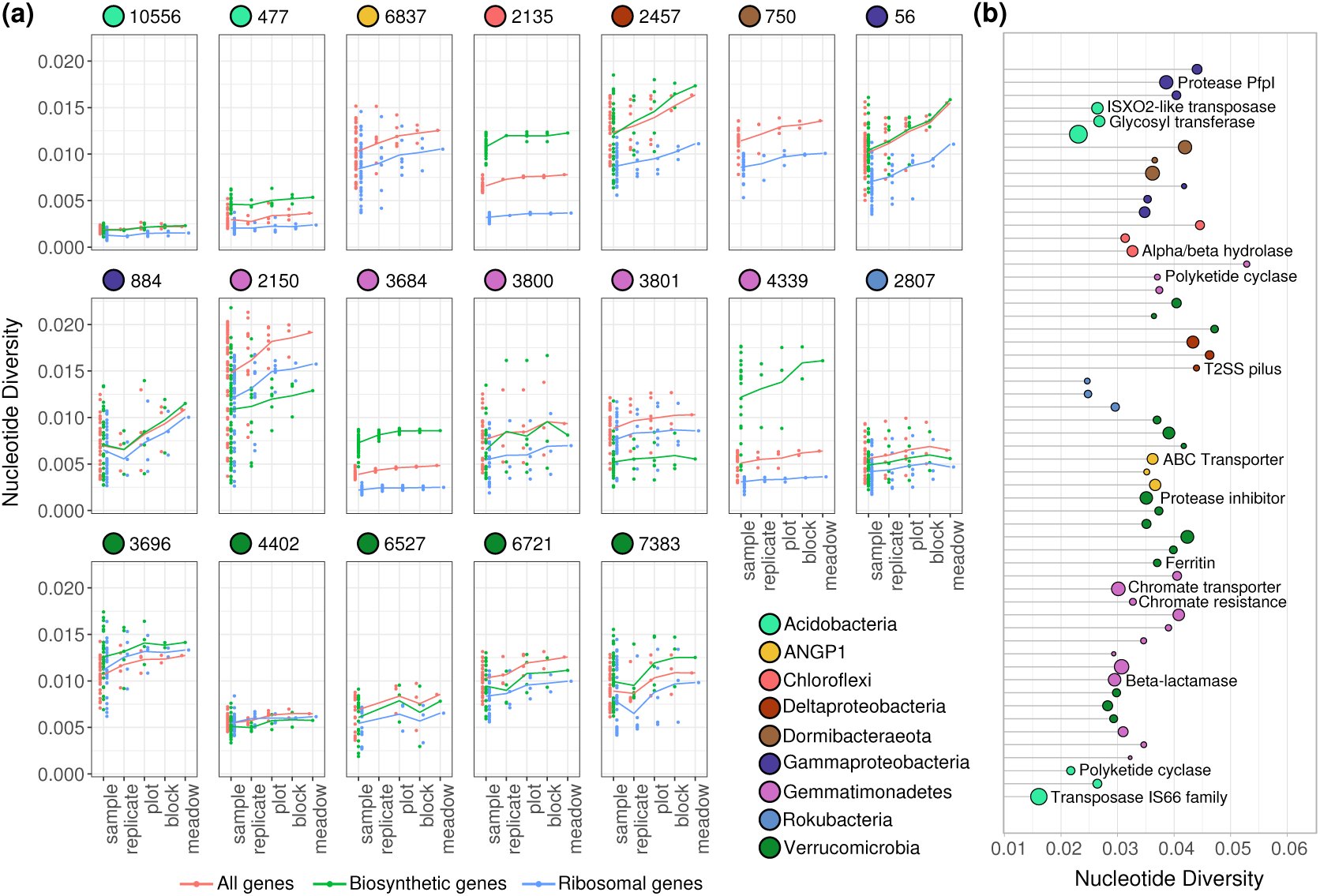
Genetic diversity of 19 highly abundant bacterial populations. **(a)** Distributions of average per-gene nucleotide diversity for each population, measured across increasing scales of sampling. Separated by all genes (red), ribosomal genes (blue), and biosynthetic genes (green). Lines connect the means of each distribution of points across scales. **(b)** Nucleotide diversity of the three most genetically diverse genes in each population. Point size scales with the gene’s length, labels indicate PFAM annotations of individual genes.

In almost all populations, the ribosomal genes consistently had lower genetic diversity than the average genes in the genome, consistent with these genes being strongly conserved and under higher purifying selection (Jordan et al. 2002) (**Fig 3a**). Conversely, biosynthetic genes involved in the production of small molecules, annotated with antiSMASH (Blin et al. 2017), were found to have significantly higher genetic diversity than the genomic average in Chloroflexi, two Gemmatimonadetes species and one Acidobacteria species (**Fig 3a**). Examining the most genetically diverse genes within each population (**Fig 3b**), we find that genes for extracellular transport were highly represented, along with polyketide cyclases, transposases, proteases, and a type 2 secretion system pilus gene. Therefore, these genes involved in extracellular uptake, sensing, and interaction may likely be experiencing local selective pressures for diversification across soil microbial species.

### Homologous recombination is common, but populations are not at panmixia

Measuring the impact of homologous recombination on the observed genetic diversity in a population can be accomplished with metagenomic data by measuring linkage disequilibrium of SNPs spanned by paired reads (Rosen et al. 2015; Lin and Kussell 2019). As two sites are further apart on the genome, the chance for a recombination event to occur between them increases, resulting in a characteristic signal known as linkage decay. Given 200 bp reads and 300-600 bp insert sizes, we could reliably assess genomic linkage of SNPs up to 800-1000 bp apart in each population. Consistent with the expectation that natural bacterial populations can undergo extensive homologous recombination, we observed the *r*^2^metric of linkage disequilibrium decay as the genomic distance between two polymorphisms increased (**Fig S2**). In the less genetically diverse populations (**Fig 4a**), there was a noticeable higher *r*^2^ of synonymous variants linked to other synonymous variants than for nonsynonymous variants linked to other nonsynonymous variants. This has been previously observed in hot-spring Cyanobacteria, and was explained as a decrease in coupling linkage for slightly deleterious nonsynonymous variants, where recombinants inheriting the doubly deleterious haplotype (of a pair of variants) are selected against (Rosen et al. 2018). In more genetically diverse populations, this ratio shifts towards nonsynonymous polymorphisms (*r*^2^_*N*_) having higher *r*^2^ than synonymous (*r*^2^_*S*_) (**Fig 4b**). One recent study reported a similar positive *r*_*N*_ *-r*_*S*_ ratio as a signature of positive balancing selection in Neisseria gonorrhoeae (Arnold et al.); for six of the 19 populations analyzed here, the genomic average *r*^2^_*N*_ /*r*^2^_*S*_ ratio was greater than 1 (**Fig 4c**), possibly also indicating recent balancing selection or admixture in this subset of populations. The increase in the *r*^2^_*N*_ */r*^2^_*S*_ ratio with nucleotide diversity (linear regression; R^2^=0.29; p=0.009) suggests a population shift from mostly slightly deleterious nonsynonymous SNPs to a higher proportion of nonsynonymous SNPs under positive selection as diversity increases.

**Figure 4.**
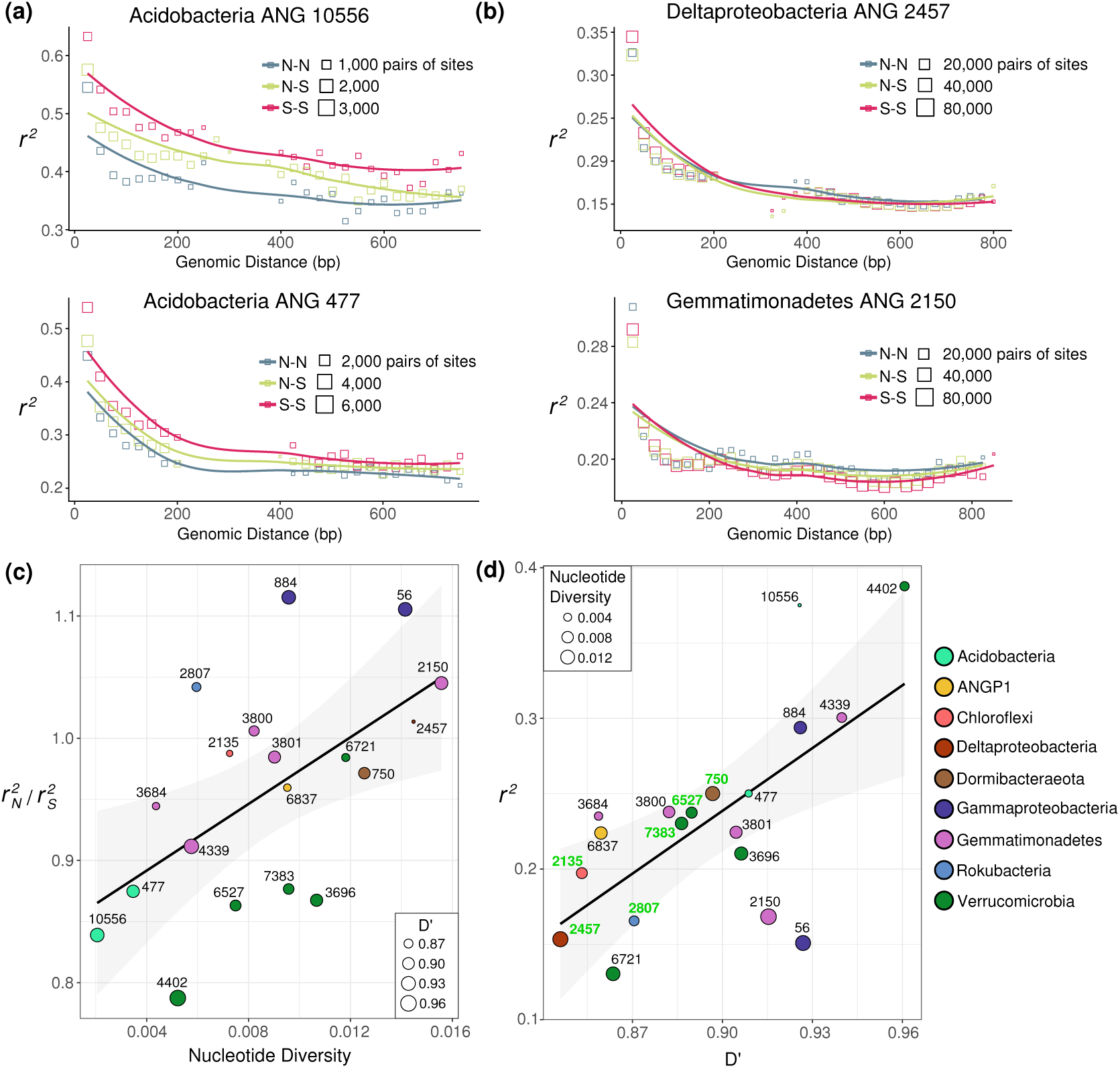
Varying rates of linkage disequilibrium within populations. **(a, b)** Linkage decay of *r*^2^ for pairs of loci within the two populations with **(a)** the lowest nucleotide diversity, and **(b)** the highest nucleotide diversity. Each square is an average of pairs of biallelic sites at that distance, with the area of the square point proportional to the number of pairs of biallelic sites that went into the mean. Haplotypes (site pairs) are binned by the predicted function of the mutations of each of the paired SNPs (nonsynonymous: N, synonymous: S). **(c)** The relationship between nucleotide diversity and *r*^2^_*N*_ */r*^2^_*S*_. The mean nucleotide diversity and the mean ratio of the linkage of nonsynonymous-nonsynonymous vs synonymous-synoymous pairs of mutations across species is shown. The size of each point represents the mean D’ value for that species. A linear regression is shown (linear regression; R^2^=0.29; p=0.009). **(d)** The relationship between mean *r*^2^ and mean D’ across the 19 bacterial populations studied. Genomes with evidence for multiple operonic competence related genes are labeled in green. A linear regression model is shown (F-statistic: 11.9, Adjusted R^2^: 0.38, p=0.003).

Although *r*^2^ is often used as a signal to identify the presence or absence of recombination, *r*^2^ values < 1 can occur with or without recombination. For example, three of four possible haplotypes (pairs of variants) for two biallelic sites could occur due to lineage divergence prior to mutation occurring at one of the sites. D’, an alternative metric of linkage equilibrium, is only <1 if all possible combinations of a pair of biallelic sites are observed (VanLiere and Rosenberg 2008), which can only occur in the presence of recombination or recurrent mutation. Generally, we found that average D’ for a population linearly correlated with mean *r*^2^ (Fig 4d). Deviation from a linear relationship between *r*^2^ and D’ (observed for three populations) could also be explained by rates of recombination being high and clonal diversity being low or vice versa. Inspection of the distribution of pairs of SNPs separated by <∼1 kbp revealed that 3 of 4 possible haplotypes is most common, but there was detection of all four possible biallelic haplotype combinations (D’ < 1), at 7% to 36% of all site pairs in each population (Fig S3). The observed frequencies of the least common of the four SNP combinations also were higher than expected based on sequencing error, although much less frequent than expected from linkage equilibrium, as evidenced by mean D’ values above 0.8. Nonetheless, the extensive appearance of D’ <1 at a significant fraction of loci, along with a signal of linkage decay with genomic distance across all populations, provides firm evidence for ongoing processes of within population homologous recombination, albeit to different degrees between organisms.

Given evidence for recent homologous recombination, we searched the genomes for genes that could confer natural competence, such as homologs of the ComEC gene with all three functional domains (Pimentel and Zhang 2018), and identified loci with additional genes involved in DNA uptake and recombination (Cassier-Chauvat et al. 2016) (**Supplementary Table S3**). We found that presence of a ComEC homolog with multiple adjacent operonic recombination-related genes was strongly associated with the lowest values of D’ (**Fig 4d**). Thus, it is likely that natural competence is a common mechanism that facilitates homologous recombination for abundant soil bacteria.

### Gene-specific selective sweeps lead to divergence of allele frequencies across the meadow

In neutrally evolving local populations, nucleotide diversity will increase monotonically with population size. Because we see no such relationship between genetic diversity and relative abundance, purely neutral growth and processes cannot explain the observed differences in genetic diversity between these populations (**Fig S4**; linear regression; p=0.88). Except for populations with the lowest genetic diversity, ratios of non-synonymous to synonymous polymorphisms for each population are consistently low. This indicates that purifying selection has eliminated slightly deleterious mutations in the populations that have accrued more genetic diversity over time (**Fig S5**; linear regression; R^2^=0.25; p=0.018). In all species except the least diverse population (an acidobacterium), non-synonymous variants also had consistently higher values of D’ than synonymous variants. Further, as genome wide D’ decreased (more recombination), the degree to which non-synonymous variants were more linked than synonymous (D’_*N*_ /D’_*S*_) increased (**Fig S6-S7**). As recombination, nonsynonymous linkage increased in comparison to synonymous linkage. This effect is consistent with stronger selection on non-synonymous variants: both purifying selection and positive selection would increase D’ for nonsynonymous SNPs.

Soil ecosystems are exceptionally heterogeneous, and environmental factors can change over millimeter distances, potentially due to changes in aboveground plant productivity, soil geochemistry, plant litter composition, and soil particulate structure (Vos et al. 2013). While it is diZcult to tease apart the effects of changing abiotic parameters over spatial scales, it is possible to examine how allele frequencies change over these scales. We calculated pairwise fixation indexes F_*ST*_ (a measure of differences in allele frequencies between two populations) for each gene between allele frequencies from the three meadow blocks for every species-group (**Fig 1b**). For most populations, mean gene F_*ST*_ values were low, <5%, consistent with dispersal of most alleles between blocks (**Fig S8**). For only a minority of populations, mean gene F_*ST*_ values were consistently >10%, indicating that there was significant geographic organization of genetic structure at most loci across the genome. Therefore, while the total variation in genome-wide major consensus alleles is often well explained by meadow geography, most individual alleles have a high chance of being found at fairly similar frequencies across the meadow.

When specific loci are characterized by significantly higher F_*ST*_ than the background average for the genome, it is characteristic of geographic location-specific selective pressures acting on that locus. To identify genomic regions of unexpectedly high F_*ST*_, we scanned over a moving 5 gene window and tested if the mean genetic diversity of that region had an F_*ST*_ greater than 2.5 standard deviations above the genomic mean (**Fig S9**). We removed genes with abnormally low coverage in either block from this analysis. To define the length of the genomic region with elevated F_*ST*_ values, we extended successful windows until the mean F_*ST*_ fell below this cutoff. This test identified a number of regions of high F_*ST*_ within some microbial genomes, despite those genomes having low average F_*ST*_ (**Fig 5a**). To further test for evidence of recent selection at these loci we looked for a statistically significant average increase in linkage and a significant change in nucleotide diversity compared to the genomic average in one or both of the blocks. We noticed that both signals of purifying selection (characterized by low N:S ratios) and a reduction in genomic coverage (potentially indicating gene loss in some portion of the population) often correlated with low nucleotide diversity in genomic regions, and based on this, caution against identifying gene-specific selective sweeps in metagenomic data based solely on a reduction in nucleotide diversity or SNP frequency. While we found many loci with unusual F_*ST*_ or strong changes in nucleotide diversity, our stringent criteria narrowed those sets down to 8 high F_*ST*_ loci with significantly increased rates of linkage compared to the genomic background (**Fig 5c**). These loci also had significant changes in nucleotide diversity within blocks when compared to their genomic averages (**Fig 5b** and **5c**). All of these loci showed decreases in genetic diversity, consistent with selective sweep events in one or multiple meadow blocks. These loci also genes had higher N:S ratios than genomic averages, possibly consistent with either recent selection acting on beneficial nonsynonymous mutations or a local accumulation of slightly deleterious nonsynonymous genetic hitchhikers.

**Figure 5.**
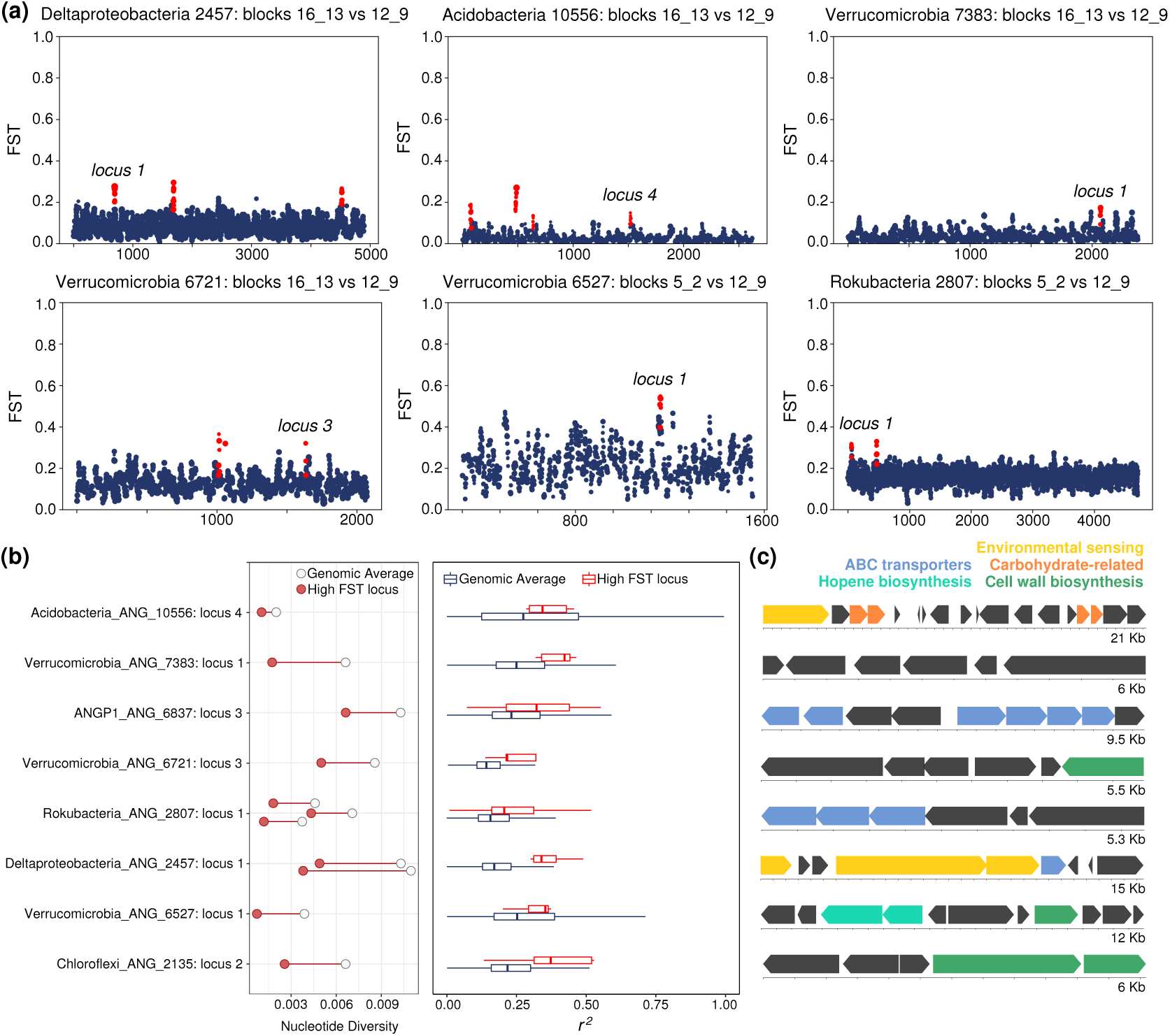
Highly differentiated genomic loci between sites within a meadow. **(a)** Values of F_*ST*_ for genes across the genomes of 6 bacterial populations. Each point is a gene, and the size of the point is determined by the number of SNPs within that gene. Plotted is the mean F_*ST*_ for that gene. Loci with significantly higher F_*ST*_ than the background are highlighted in red, and labeled by locus numbers used in part (b). **(b)** Left: Nucleotide diversity at highly differentiated loci (red circle) compared to the average (empty circle) for each population. Right: The extent of linkage disequilibrium at highly differentiated loci (red) compared to the genomic average (black) for each population. **(c)** Gene diagrams and annotations of highly differentiated loci (genome and loci identities given in panel b). Each block indicates an open reading frame, and blocks are colored by a subset of predicted functions.

The loci with evidence of recent differential selection across the meadow commonly contained transporter genes (**Fig 5c, Supplementary Table S4**), which could indicate selective pressures for uptake of different compounds between sites. A Verrucomicrobia population also showed evidence of a selective sweep occurring at a putative hopene biosynthesis operon, often involved in regulating membrane stability. Within a Deltaproteobacteria population, a highly differentiated locus encoded for numerous genes involved in two-component systems and histidine kinases, potentially related to environmental sensing and response. Taken together, these multiple genomic signals suggest that gene-specific selection drives differences in population genetic structure across meadow soils.

## Conclusions

Developing a cohesive picture of the genetic structures of soil bacterial populations is crucial for understanding the evolution and distribution of genes that can play critical ecosystem and niche-specific functions. Doing so has been diZcult because the most abundant soil bacteria are diZcult to cultivate, and when cultivation is possible, it is uncertain if microbial isolates are truly random samples of a population, considering the intense selective pressure put on cells during the isolation process. Here, we show that recent advances in sequencing technologies and software tools enable the monitoring of heterogeneity within genomically-defined bacterial populations needed to address these questions in soil.

Population genetic diversity in abundant soil microbes was higher than that reported for populations in other environments, mirroring the high species diversity in soils. It is possible that the spatial and biogeochemical complexity of soils promotes niche formation that leads to both increased species diversity and in turn more genetic diversity within those species. This possibility has direct implications for reductionist approaches in soil microbial ecology using synthetic communities of clonal isolates. Synthetic communities are qualitatively different from actual soil microbial communities: in real soil communities, each species contains allelic and gene content diversity that is highly variable, while recombination generates a continuum of genetic distributions of alleles. Therefore, population-level diversity in natural soil communities can expand the available biotic niches into which genotypic variants adapt, which can produce qualitatively different community dynamics and outcomes than synthetic mixtures of genetically identical strains, even if species composition is consistent.

The results of this study suggest that recombination and gene-specific selection are important modes of evolution in soil microbes. Even in a single meadow we predicted that there are likely thousands of combinatorial genetic mixtures for each species, with recombination resulting in no easily measurable numbers of irreducible strain-like lineages. Thus, the dynamics of dominant soil bacterial populations may be partially described using ‘quasi-sexual’ models (Rosen et al. 2015; Garud et al. 2019). Although we only observed the importance of these dynamics in structuring populations at the 1-10 meter scale, it is still uncertain how evolutionary dynamics in soil bacterial populations develop on other spatial and temporal scales. We conclude that future work on soil microbial ecology would benefit by considering the role of unlinked and substantial allelic diversity within species in shaping local gene content and allele frequencies.

## Materials and Methods

### Sampling, genome sequencing, and metagenomic assembly

The sampling scheme, local soil characteristics, and study design that were utilized in this analysis have been previously described (Diamond et al. 2019). The 60 soil cores were sampled within the 6 plots, each 10 meters in diameter, over a period of two months before and following autumn rainfall. Every plot was directly adjacent to a paired plot (forming a “block”). Briefly, DNA was extracted from 10 g of soil for each sample using the PowerMax Soil DNA isolation kit (MoBio Laboratories) from individual soil cores. Metagenomic libraries were prepared and sequenced with 2×250 paired read sequencing on the Illumina HiSeq2500 platform at the Joint Genome Institute. Reads were quality filtered to a maximimum 200 bp in length using BBduk, and the target inter-read spacing was 500 bp. Metagenomes were assembled using IDBA_UD (Peng et al. 2012) and individual genomes were binned using differential coverage beginning and a suite of metagenomic binners as previously described (Diamond et al. 2019).

### Genome dereplication, filtering, and comparison

The 10,538 genome bins previously described were dereplicated using dRep (Olm et al. 2017) with the secondary clustering threshold -sa 0.97. CheckM (Parks et al. 2015) was used to evaluate genome completeness and contamination. For this analysis, we then analyzed species with at least 12 dereplicated genomes that were estimated to be >80% complete with <10% contamination, independently assembled and binned out of different samples. Open reading frames were predicted using Prodigal (Hyatt et al. 2010) and annotated using (1) USEARCH against UniProt (UniProt Consortium 2018), Uniref90, and KEGG (Kanehisa 2000), and (2) HMM-based annotation of proteins using PFAM (Finn 2005), and (3) antiSMASH (Blin et al. 2017) for biosynthetic gene prediction. PERMANOVA tests for association of ANI matrices with environmental data were run using the vegan package (Dixon 2003) in R (R Core Team 2018).

Possible contamination was stringently removed from representative genomes using the assembled replicate genomes for each species population. The pan-genomic analysis pipeline Roary (Page et al. 2015) was run with default settings on the set of genomes for each species. Contigs with at least 50% of their gene products found in less than 25% of each genome set were then discarded as potential contamination (generally fewer than 20 contigs, often small, were removed per genome). Therefore, the final set of contigs used in this analysis contained only contigs that reliably assembled and binned independently for a species in multiple samples. Organism relative abundances were calculated using the total number of reads that mapped to each genome in each sample and dividing by the total number of reads per sample.

### Read mapping, SNP calling, and nucleotide diversity

All metagenomic reads were mapped using Bowtie 2 (Langmead and Salzberg 2012) with default parameters except for the insert size parameter -X 1000 to an indexed database of the 664 dereplicated genomes obtained from the environment. Read filtering of the resulting BAM files was performed using a custom script, filter_reads.py, available at the link under code availability. Mapping files were then filtered for reads that meet the following criteria: (1) both reads in a pair map within 1500 bp of each other to the same scaffold (counting end to end insert size), (2) the combined read pair maps with a percent identity of at least 96% to the reference, (3) at least one of the read pairs has a mapq score > 1, indicating that this is a uniquely best mapping for this read pair in the index. Manual inspection of representative mapping files indicated that these filters greatly decreased rates of read mismapping and erroneous read pileups.

For all population diversity metrics, we used all filtered cross-sample read mappings that passed a cutoff of at least 50% of the genome being covered with at least 5× coverage. 586 out of 1140 sample genome comparisons (19 genomes*60 samples) passed this minimum requirement. Base pairs of reads with less than Phred scores of 30 were not used in SNP or linkage analyses. Nucleotide diversity was calculated on all genomic positions with at least 5x coverage within each sample. Sample read mappings were pooled by replicates, by plot of origin, by block and origin, and finally by all samples in the meadow, and nucleotide diversity was recalculated on each pooled set of samples. For downstream analyses, nucleotide diversity for all samples (meadow-wide, **Fig 3**) and nucleotide diversity within each of the three sampling blocks (block-wide, **Fig 5**) was analyzed.

SNPs were called on the combined meadow-wide population set of filtered reads. We constructed a simple null model based on an error rate of 0.01% (the Phred 30 cutoff) and simulated simple sampling with replacement to construct estimated rates of erroneous SNP counts at pileups of varying coverages. Positions with an alternative allele that occurred with counts that had a false positive rate of ∼10^-6^ with the given coverage of that site in the null model and a minimum allele frequency (MAF) of 5% were called as SNPs. Because meadow-wide coverages were at least 224× (and ranged up to 908x) the MAF cutoff alone would likely be a stringent cutoff for Phred 30 error rates. SNPs were assigned as synonymous or non-synonymous using a custom BioPython script and the gene calls annotated by Prodigal.

### Linkage disequilibrium, FST, and tests for selection

Linkage was calculated for all pairs of segregating sites that were spanned by at least 30 read pairs with high quality base pair data. *r*^2^ and D’ linkages were calculated using formulas described by (VanLiere and Rosenberg 2008). F_*ST*_ was calculated on sites segregating across both blocks being compared (for all three block comparisons) using the Hudson method (Hudson et al. 1992) as recommended by (Bhatia et al. 2013), as implemented in the scikit-allel package (Miles et al. 2019). A site had to have a coverage of at least 20x in each block in order to calculate F_*ST*_, and genes which had coverages in a block outside of the range of two standard deviations were excluded from the analysis. A ratio of averages was then used to determine mean F_*ST*_ for each gene. A two-sample Wilcoxon test was used to determine if average linkage of highly differentiated loci differed from the genomic average for each species, and two-sample t-tests in R (R Core Team 2018) were used to determine if average nucleotide diversity of highly differentiated loci differed from the genomic average. Both sets of tests were corrected for multiple hypotheses using the Benjamini-Hochberg (Benjamini and Hochberg 1995) method.

## Code and data availability

The published genomes and raw sequencing reads for this study are available under NCBI BioProject number PRJNA449266. Reproducible notebooks, analyses and additional data tables used in this analysis are available at https://github.com/alexcritschristoph/soil_popgen/. Large data tables and genome sequences are available at: https://figshare.com/projects/Soil_bacterial_populations_are_shaped_by_recombination_and_gene-specific_selection_across_a_meadow/65789

## Acknowledgments

The authors thank S. Spaulding for assistance with fieldwork and B. Good for helpful discussions on data analysis and theory. The authors thank K.B. Suttle for establishing the field site’s grassland plots and maintaining experimental precipitation regimes and for the aerial photograph in Figure 1. Sampling was performed at the Angelo Coast Range Reserve and made possible by the University of California Natural Reserve System and the National Science Foundation CZP EAR-1331940 for the Eel River Critical Zone Observatory. Sequencing was carried out under a Community Sequencing Project at the Joint Genome Institute. Funding was provided by the OZce of Science, OZce of Biological and Environmental Research, of the US Department of Energy (grant DOE-SC10010566).

## Supplemental Figures

**Supplementary Figure S1.**
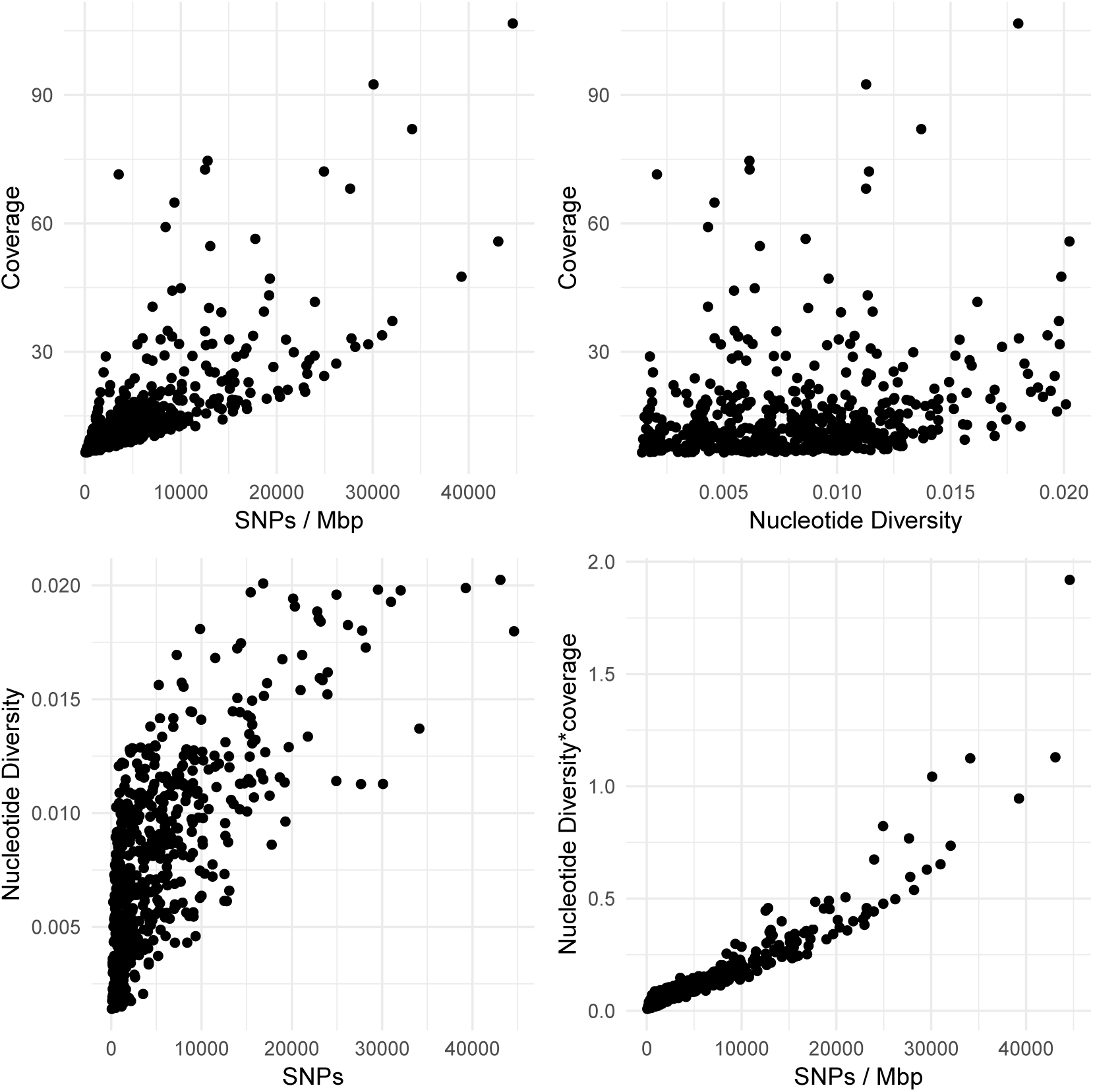
The impact of coverage on the number of SNPs called in each sample. Each point is a sample-genome mapping pairing, with the number of SNPs called per sample plotted against the mean coverage of the genome in that sample, the mean nucleotide diversity of the genome in that sample, and the mean nucleotide diversity times the coverage, demonstrating that at low coverages the number of SNPs called is a function of nucleotide diversity and coverage.

**Supplementary Figure S2.**
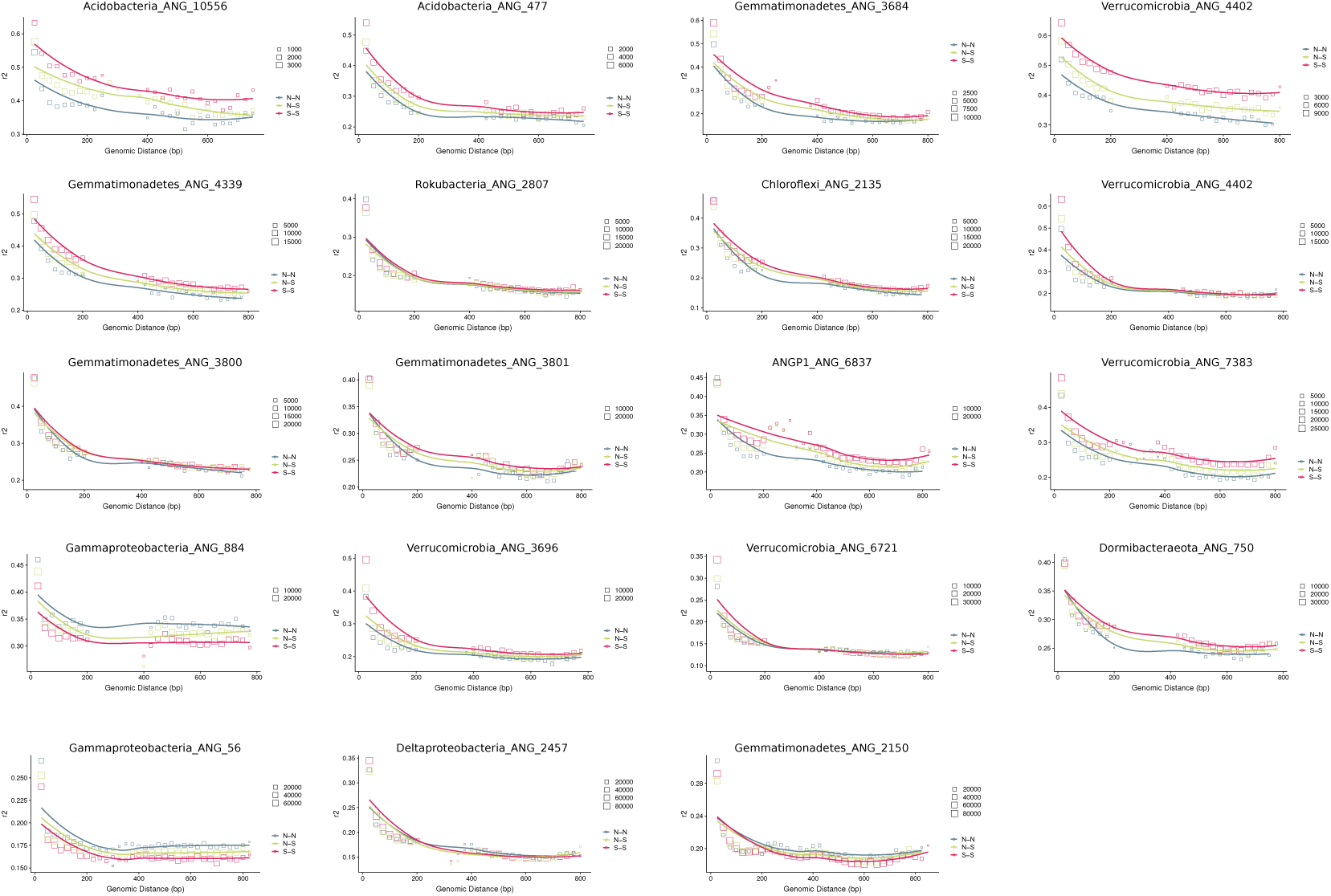
Linkage disequilibrium decay over genomic distance for all 19 species in this study. Each block point represents a mean of linkage for SNPs at that genomic distance, divided into non synonymous-nonsynonymous linkages, nonsynonymous-synonymous linkages, and synonymous-synonymous linkages. The size of each block point represents the number of SNPs that went into calculating the mean.

**Supplementary Figure S3.**
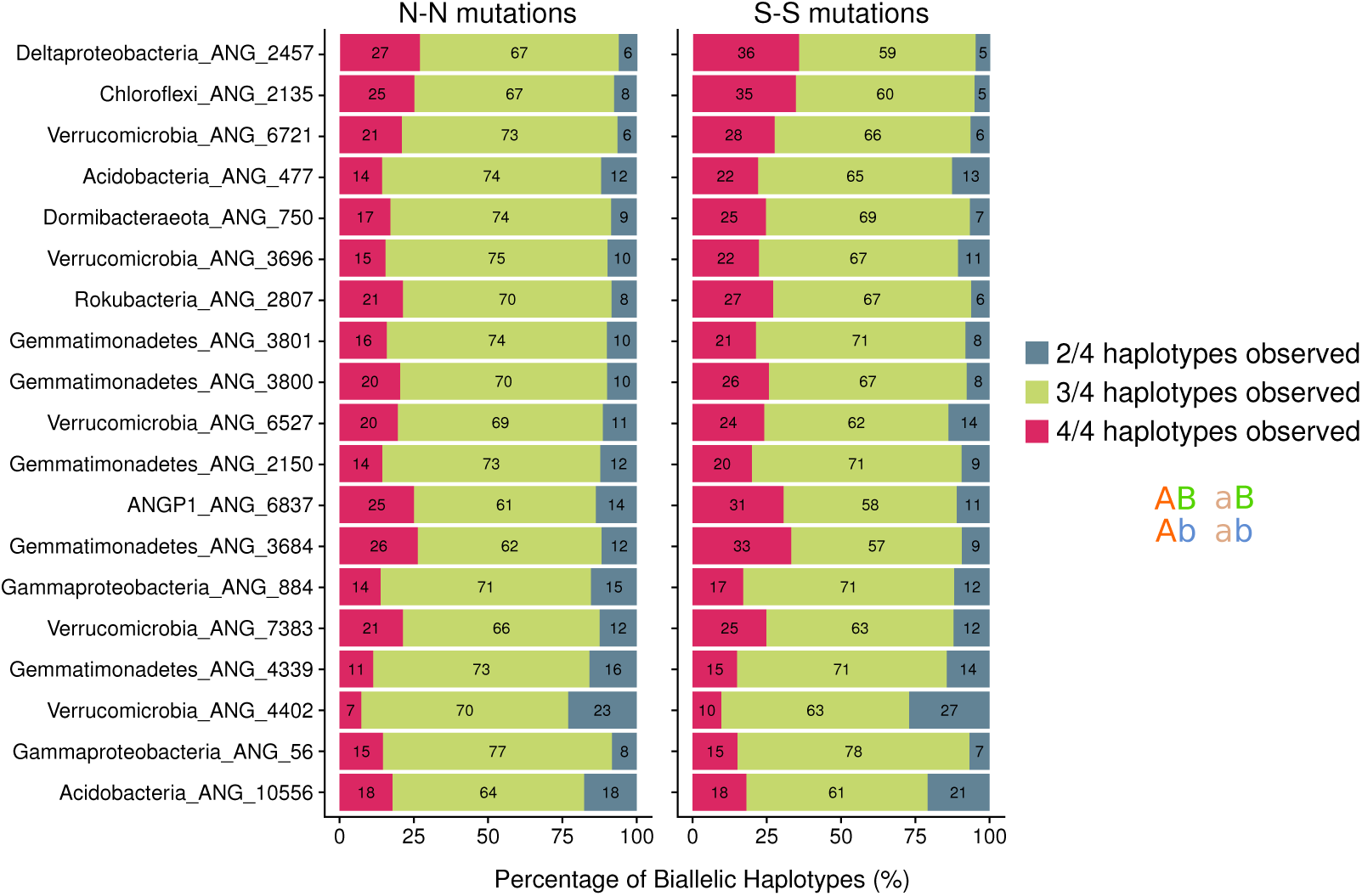
Biallelic haplotype counts for each species. The percentage of haplotype counts for each pair of biallelic sites within ∼1 Kb of each other is shown for both nonsynonymous-nonsynonymous and synonymous-synoymous pairs of segregating sites.

**Supplementary Figure S4.**
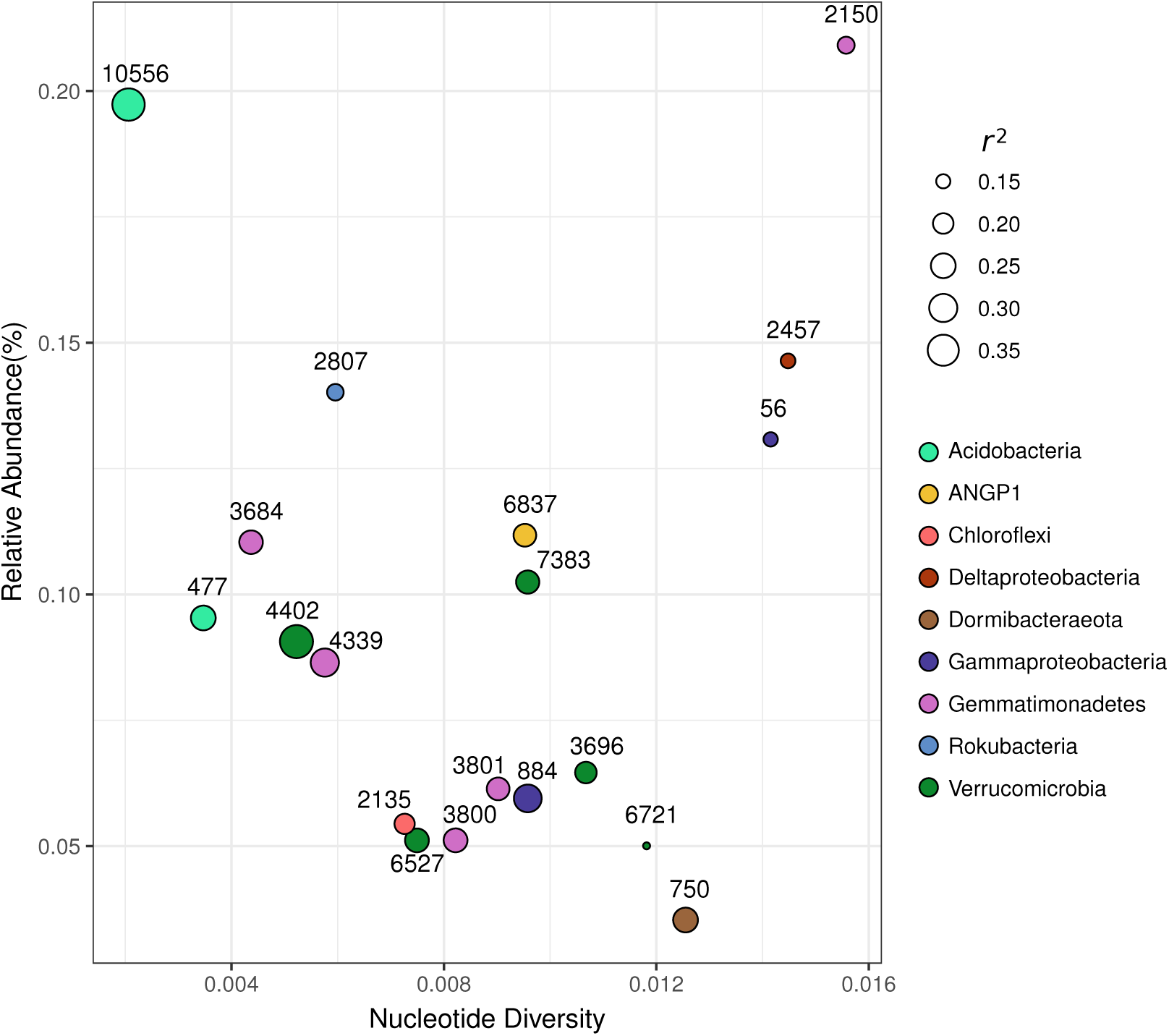
Relationship between nucleotide diversity and relative abundance across species.

**Supplementary Figure S5.**
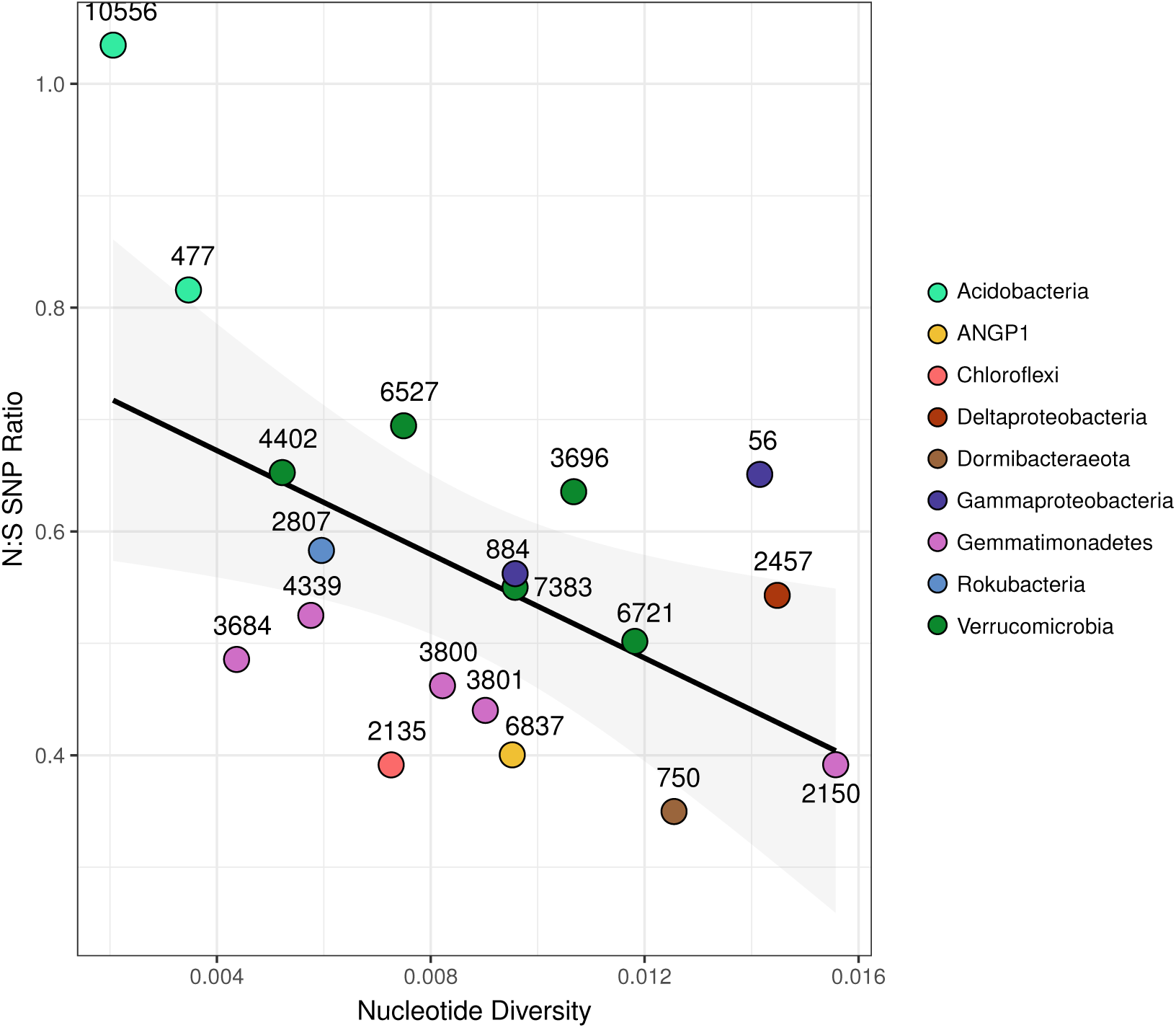
Relationship between nucleotide diversity and the N:S SNP ratio across species. A linear regression is shown (R2=0.25; p=0.018).

**Supplementary Figure S6.**
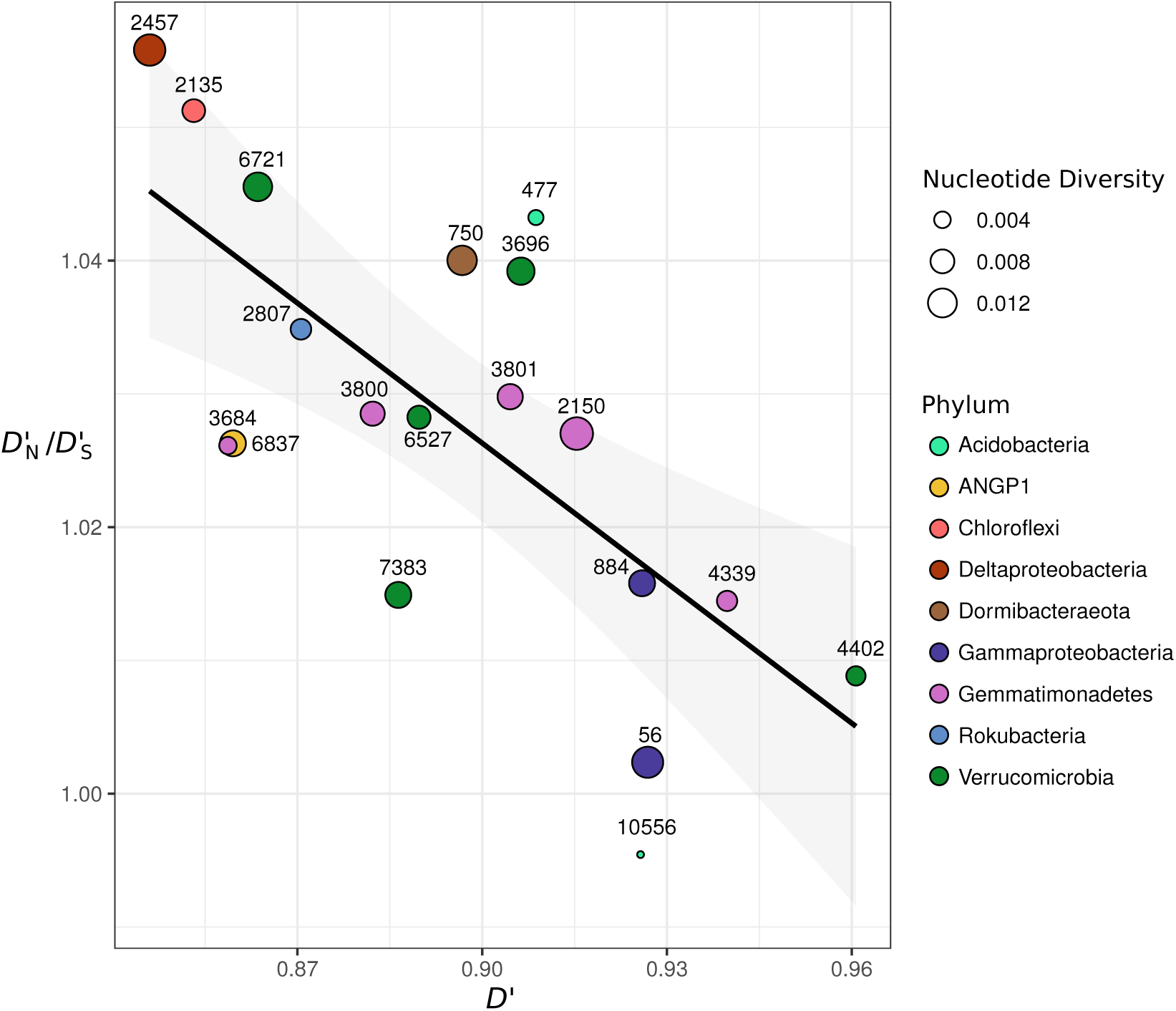
Relationship between D’ and D’_*N*_ /D’_*S*_. The ratio of non-synonymous polymorphism linkage to synonymous polymorphism linkage (D’N/D’S) versus mean D’ for all bacterial populations studied. A linear regression model is shown (F-statistic: 15.6; Adjusted R-squared: 0.45, p=0.001).

**Supplementary Figure S7.**
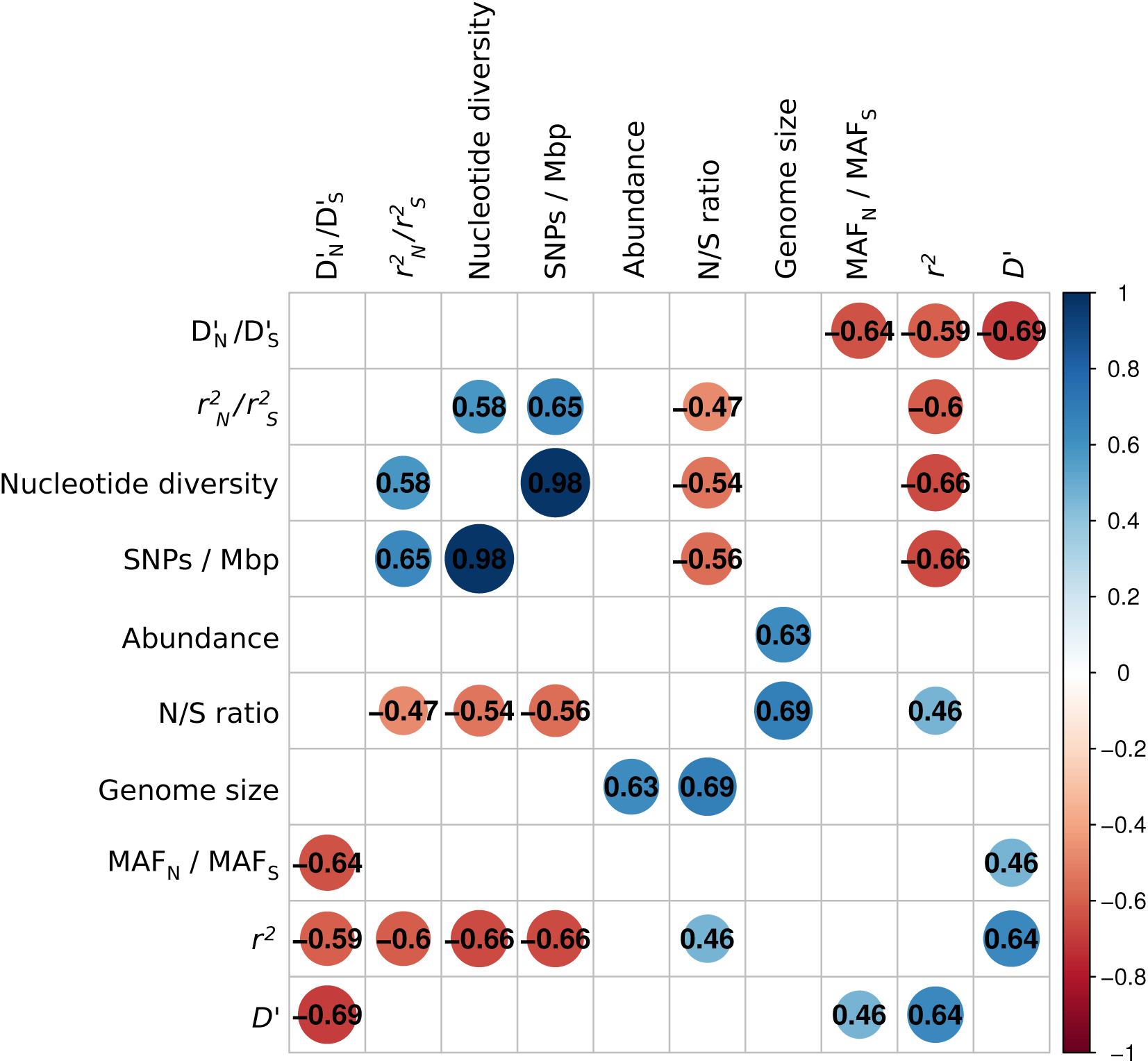
Cross-population correlations between means of all population genetics summary statistics in this study. In each box, the pearson correlation coeZcient between two metrics is shown. Only correlation coeZcients with a p-value <0.01 are shown.

**Supplementary Figure S8.**
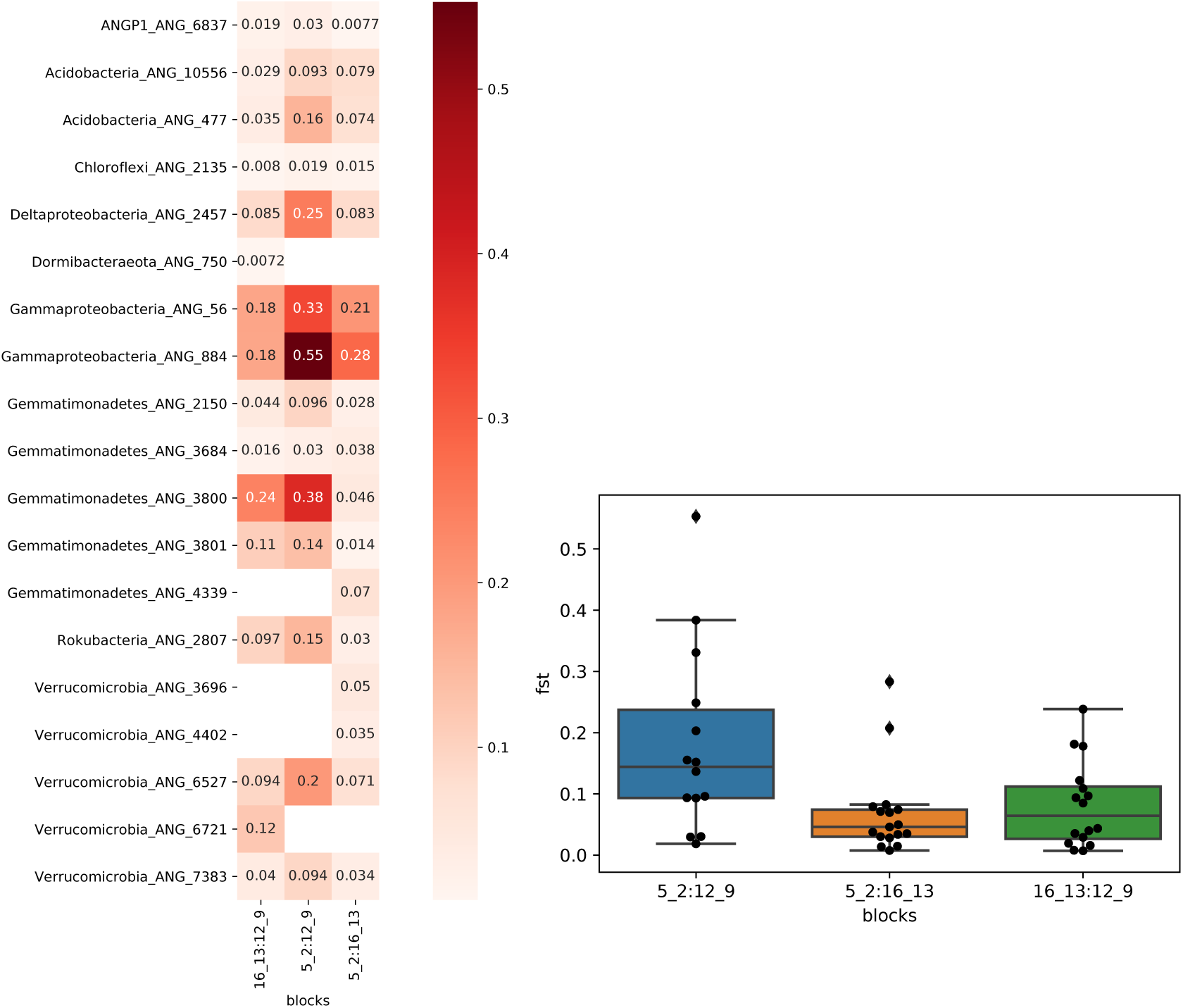
Mean FST values for every species across the meadow. The mean per-gene FST values for each species between each of the three block comparisons is shown.

**Supplementary Figure S9.**
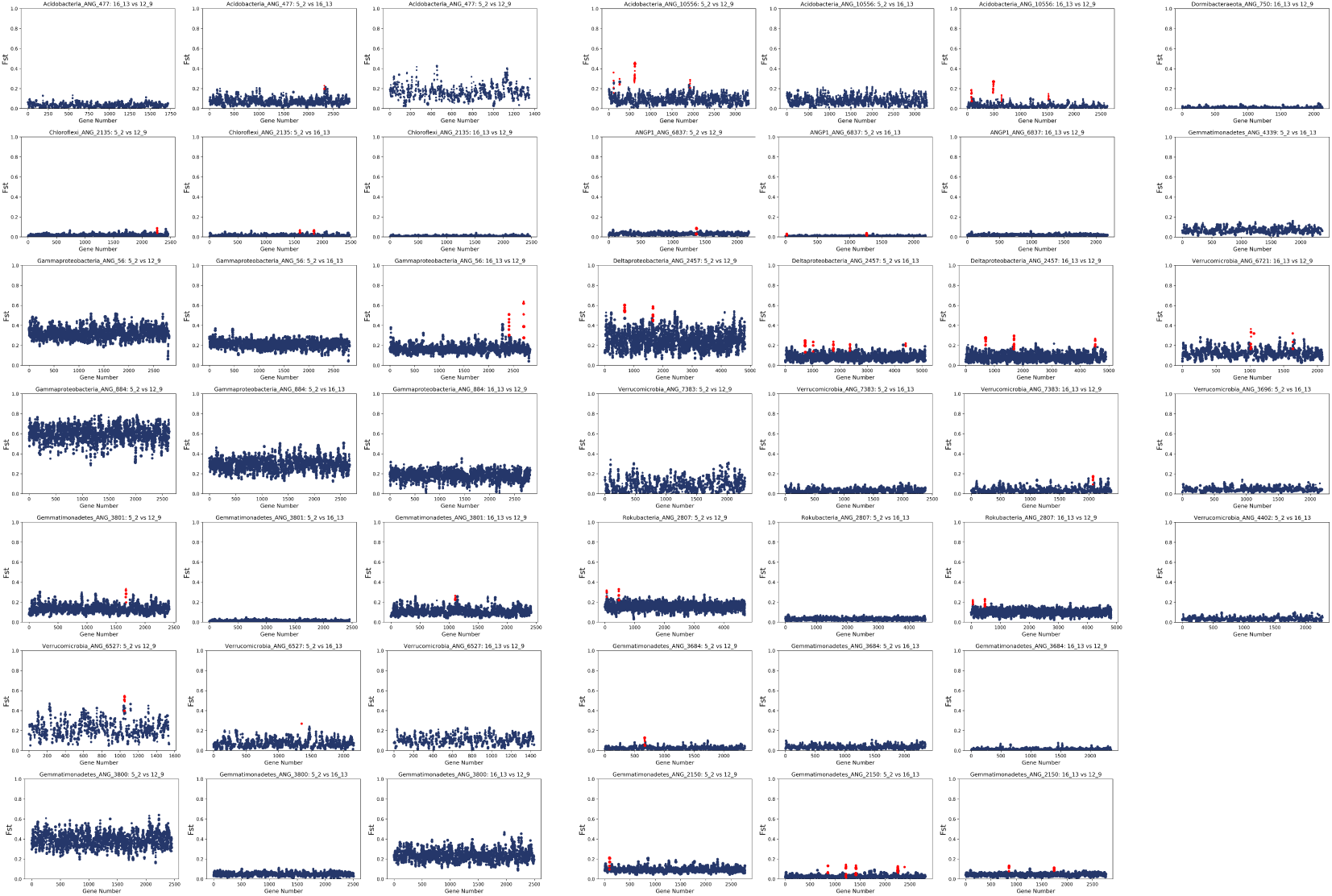
Gene FST along the genome for every species across the meadow. Per-gene FST values are plotted in ORF order on contigs. The title of each plot shows the species name and the block vs block comparison values being plotted. Each point is a gene, and the size of the point increases with the number of segregating sites in that each. Genes are colored in red if they are part of a locus with mean FST greater than 2.5 times the standard deviation of the genomic average.

**Supplementary Table S1** Completeness and contamination statistics for all population replicate genomes included in this study.

**Supplementary Table S2** Key population genetics summary statistics in this study.

**Supplementary Table S3** ComEC annotations for each species.

**Supplementary Table S4** Annotations of genes within highly differentiated loci.

